# Role of Dienelactone Hydrolases in PET Biodegradation by Flavobacteria *Maribacter dokdonensis* and *Arenibacter palladensis*

**DOI:** 10.1101/2025.08.25.672177

**Authors:** Ly T.T. Trinh, Ifey Alio, Pablo Perez-Garcia, Sabine Keuter, Robert Dierkes, Lena Preuss, Christel Vollstedt, Wolfgang R. Streit

## Abstract

Dienelactone hydrolases (DLHs, EC 3.1.1.45) are enzymes that play a crucial role in the breakdown of cyclic esters and some have been found to act on substrates such as terephthalate esters, which are monomers of polyethylene terephthalate (PET). In the current study, we show that bacteria affiliated with the Bacteroidota (class Flavobacteria) harbor DLHs acting on PET foil and powder. We report on the isolation of two marine bacterial strains, *Arenibacter palladensis* UHH-Hm9b and *Maribacter dokdonensis* UHH-5R5, forming biofilms on PET foil and releasing µM amounts of terephthalic acid after 5-7 days. Genome sequencing and functional analyses identified two secreted DHLs designated PET93 and PET 94 involved in PET degradation. While their predicted active sites and substrates binding pockets were identical to previously published PETases, both enzymes differed largely in their structural features from known PETases and represent novel scaffolds. Further they lacked the typical porC-domain of the known PETases from the Flavobacteria. Biochemical characterization of the two recombinant enzymes confirmed activity on PET, the primary degradation products Bis(2-Hydroxyethyl) terephthalate (BHET) and Mono-(Hydroxyethyl) terephthalate (MHET). These are the first DLHs to be reported being active on plastics and our findings indicate that Flavobacteria harbor an unexpectedly wide range of PET-active promiscuous enzymes.

**IMPORTANCE:** Global plastics pollution is a major environmental challenge, and we still have limited knowledge of marine microbiota involved in possible remediation. Our research shows that marine Flavobacteria harbor the potential for PET degradation using dienelactone hydrolases (DLHs, EC 3.1.1.45). The widespread distribution of these microorganisms and the notion that these enzymes are secreted may imply a possible role in marine PET remediation.

## INTRODUCTION

Recent estimates indicate that at least 8-10 million tons of plastic waste are introduced into the ocean annually (1–4). While most of the plastic initially exists as larger floating fragments, exposure to weathering factors such as sunlight, wind and water currents gradually breaks it down into smaller particles, from micro to nano sizes which are smaller than 5 mm in diameter and more spreadable in air, water and soil (5, 6). These particles, along with the chemicals they contain, are believed to negatively impact ecological systems and biodiversity across all environments (7–9). Polyethyleneterephthalate (PET), a widely used plastic polymer, has been accumulating in the oceans over the past decades, and we have only a limited understanding of and to which extent this polymer can be degraded by microorganisms. While PET is one of the commodity polymers that can be enzymatically degraded under laboratory and industrial scale conditions, there is little knowledge on the microbiota potentially involved in marine PET degradation. Today 125 PET microbial hydrolases (wildtype enzymes) have been described (for a complete list see PAZy database, www.pazy.eu, accessed on 22.07.25) (10).

It is currently known that PET is in general degraded by promiscuous and secreted hydrolases affiliated with the esterases enzyme class (EC 3.1.1-.) They are either designated carboxylesterases (EC 3.1.1.1), poly(ethylene terephthalate) hydrolases (EC 3.1.1.101), lipases (EC 3.1.1.3) or cutinases (EC 3.1.1.74). All have a relatively wide substrate range and act mostly unspecific on the polymer (11–13). Notably, among the currently known and functional PETases no dienelactone hydolases DLHs (3.1.1.45) are listed.

Recent research from our laboratory identified PET degrading enzymes, PET27 and PET30, derived from the phylum of the marine Bacteroidota (42). The two enzymes were unique as they were secreted using the T9SS. Therefore, both enzymes harbored a porC and N-terminal secretion signal. While the overall turnover rates of the enzymes were rather low, they were active at low temperatures. The Bacteroidota phylum represents a major evolutionary lineage within the domain Bacteria and is one of the dominant components of the marine microbiomes. This phylum is particularly notable for its ability to degrade a wide range of natural polymers through multi-enzyme complexes, making them a valuable source of biologically active compounds (14). It is a highly diverse group of gram-negative bacteria that encompasses at least four distinct classes: Bacteroidia, Flavobacteria, Sphingobacteria and Cytophagia (15). Among them, Flavobacteria is the largest and most widely distributed class of Bacteroidota which are known for their surface-active properties and are frequently found associated with marine algae and sponges (16, 17). Flavobacteria are also one of the dominant classes on floating plastic particles such as LDPE, PP and PET and major contributors to marine polysaccharide degradation (18–20).

In this study, we provide strong evidence that the Flavobacterial isolates, *Maribacter dokdonensis* (UHH-5R5) and *Arenibacter palladensis* (UHH-Hm9b), code for active but promiscuous PET hydrolases, PET93 and PET94, enabling PET degradation while growing in biofilms on PET-foils. The two novel enzymes are the first DLHs (3.1.1.45) reported to be active on this polymer. Altogether our findings indicate that the Bacteroidota harbor an unexpectedly wide range of PET-active enzymes potentially involved in polymer degradation in the marine environment.

## RESULTS

### Enriching for PET-active enzymes affiliated with the Bacteroidota

To expand the functional diversity of PET-active enzymes within the phylum Bacteroidota we enriched for aerobic Flavobacterial species from environmental samples using either BMB or R2A culture medium supplemented with PET powder as described in the Materials and Methods section. The different enrichment cultures resulted in a total of 19 distinct isolates as summarized in Table 1. Among them, four isolates were included that had been obtained in previous studies from our laboratory. The phylogenetic affiliation of the isolates was initially verified using 16S rDNA gene analysis. All isolates were affiliated with the phylum of the Bacteroidota and the majority were affiliated with the class Flavobacteria. As a further initial screening to assess if any of these isolates indeed was able to secret PET-active hydrolases (I.e. esterases E.C. 3.1.1.1) the isolates were tested for their capability to hydrolyze typical model substrates commonly used to screen for PET-active bacteria (e.g. TBT, PCL and BHET) on agar plates. Halo formation was used as an indicator for the production and secretion of active hydrolases. Among all tested isolates, only two, UHH-5R5 and UHH-Hm9b, produced visible halos and were able to hydrolyze BHET while only UHH-5R5 acted on both BHET and PCL (Figure 1A). Using 16SrDNA analyses, we confirmed that UHH-5R5 and UHH-Hm9b phylogenetically cluster closely together and form a distinct branch within the Bacteroidota clade and belong to the class of Flavobacteria (Figure 1B). Strain UHH-Hm9b is affiliated with the species *Arenibacter palladensis* while the UHH-5R5 isolate is affiliated with the species *Maribacter dokdonensis* (Table 1).

**Figure 1.**
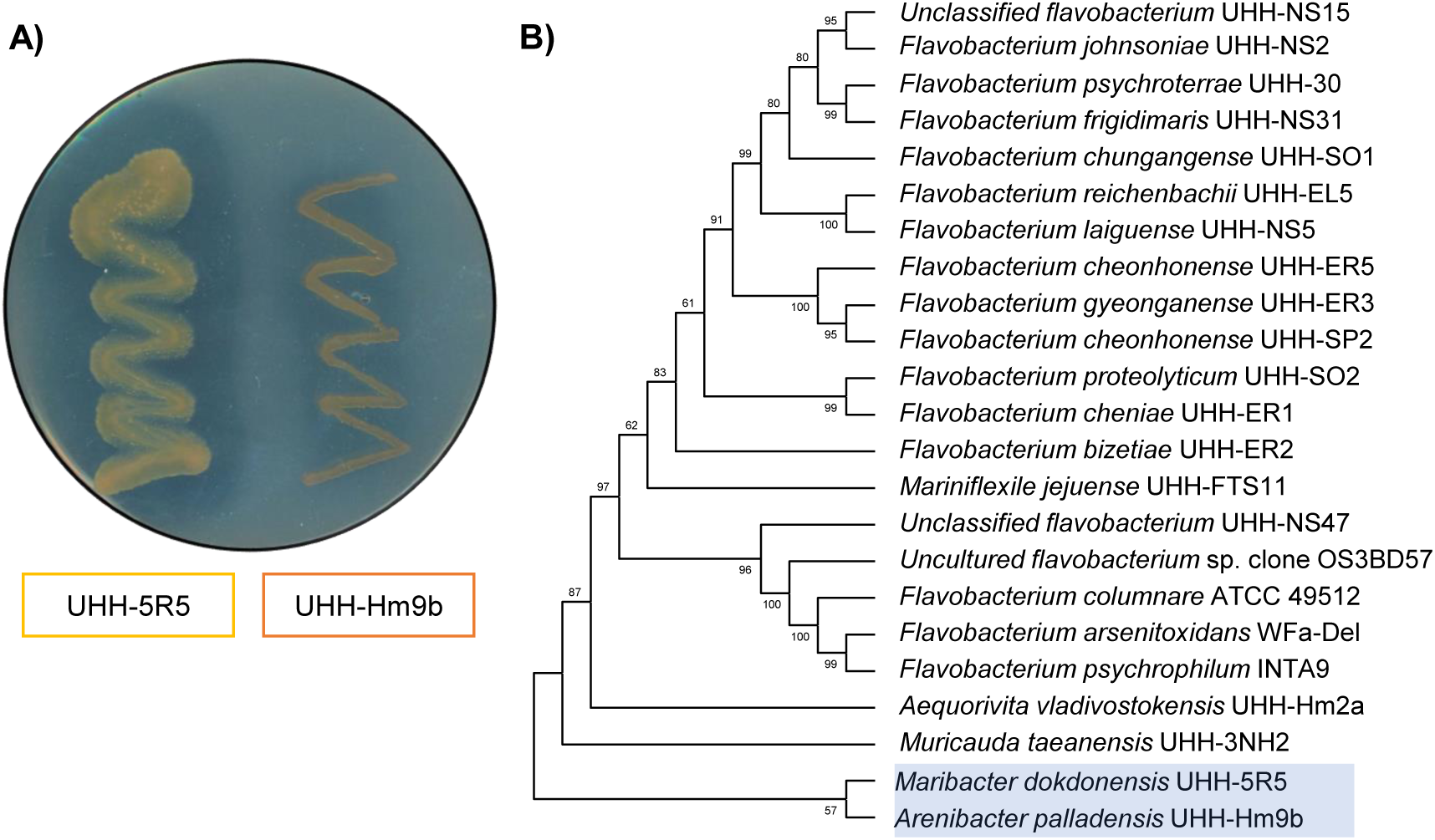
**(A)** Visualization of BHET-hydrolyzing activity by the isolates: *Maribacter dokdonensis* UHH-5R5 and *Arenibacter palladensis* UHH-Hm9b. Clear zones (halos) formed around bacterial colonies grown on BHET agar plates indicate enzymatic hydrolysis of the substrate. Plates were incubated at 28 °C for 5 days. **(B)** Phylogenetic tree based on 16S rDNA gene sequences of the two newly isolated strains (highlighted in blue) and other members of the Bacteroidota. The tree was constructed using MEGAX and the number displayed at nodes are the bootstrap values based on 1,000 replications.

**Table 1:**
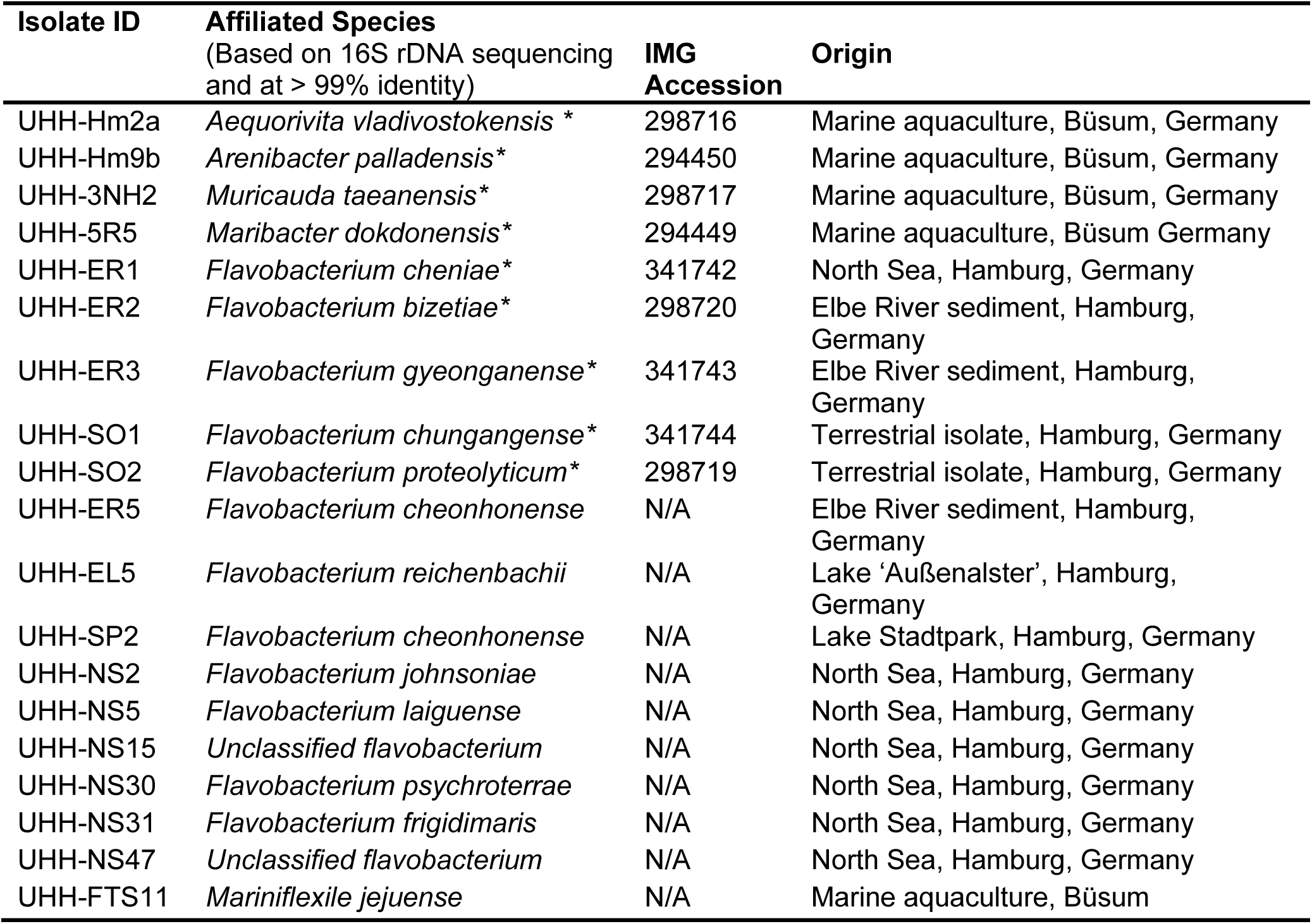
Flavobacteriaceae isolates enriched in this work using PET powder as substrate. Sequenced genomes were marked with an asterisk (*)

### *Maribacter dokdonensis* UHH-5R5 and *Arenibacter palladensis* UHH-Hm9b form dense biofilms on PET foil and release TPA

Intrigued by the above-made observations, we asked if both isolates would grow on PET foil, form biofilms, and release small amount of TPA (terephthalic acid) as one of the primary PET degradation products. To assess if UHH-5R5 and UHH-Hm9b degrade PET foil under biofilm conditions, we inoculated both wild type strains in BMB medium with PET foil as a possible substrate and surface to form biofilms on. The attachment and biofilm formation of UHH-5R5 and UHH-Hm9b on PET were observed over a seven-day incubation period using laser scanning microscopy and potential PET degradation was analyzed using UHPLC to detect the released TPA.

Initially, only a few individual cells of both strains were sparsely adhered to the PET surface, with no visible biofilm structures. However, on day 2, thin biofilm layers formed, accompanied by an increased number of attached cells (Figure 2). The 2-day old biofilms of UHH-5R5 and UHH-Hm9b exhibited an average thickness of 5.9±1.04 µm and 8.4±1.14 µm, respectively, corresponding to approximately 3–5 cell layers on foils. An increase in biofilm development was observed at day 3, with biofilm thickness reaching 6.7±2.1 µm (UHH-5R5) and 12±3.2 µm (UHH-Hm9b). At day 6, both strains developed well-structured, multilayered biofilms, reaching thicknesses of 14.5±1.8 µm (UHH-5R5) and 13.4±1.2 µm (UHH-Hm9b). Interestingly, while the biofilm thickness of UHH-5R5 decreased slightly to 11.3±2.5 µm by day 7, that of UHH-Hm9b remained stable at 15.1±3.1 µm (Supplementary Table S2).

**Figure 2.**
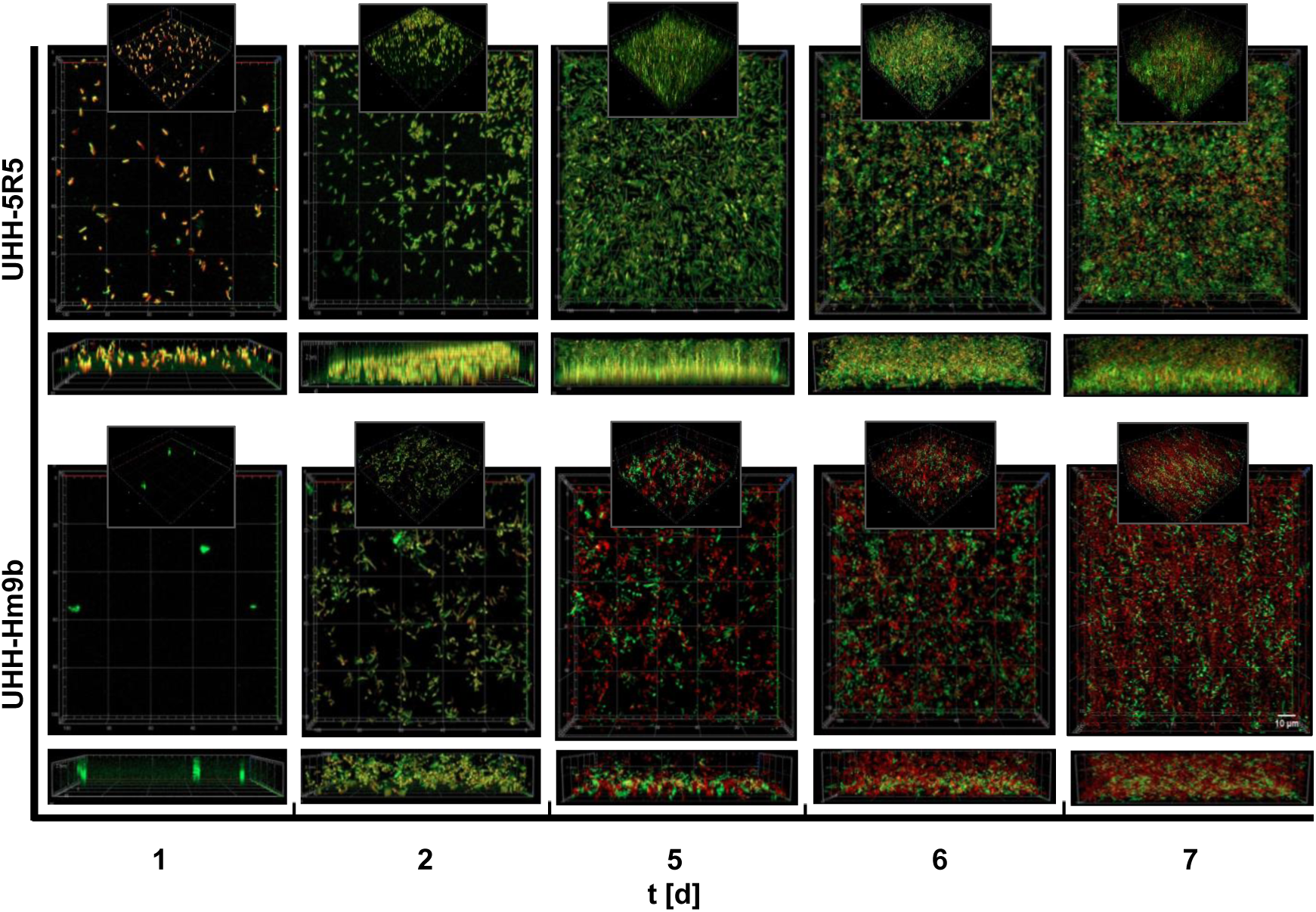
Fluorescence microscopy images showing the development of biofilms formed by the Bacteroidota isolates UHH-5R5 and UHH-Hm9b on PET foil over a 7-day incubation period. Biofilms were visualized using Live/Dead staining, where live cells fluoresce green (SYTO 9) and dead cells fluoresce red (Propidium iodide). Images were captured on days 1, 2, 5, 6, and 7 using a confocal laser scanning microscope (CLSM) with a 63× oil immersion objective. For each sample at least three different positions were observed and all images here are representative of three independent biological replicates.

To detect possible PET degradation products, we collected and concentrated supernatants from 3-, 5- and 7-day old biofilms grown on PET foil. UHPLC analysis revealed that strain UHH-5R5 released approximately 100-320 µM of TPA in the concentrated supernatants, equivalent to approximately 6-19 µM in the original samples (corresponds to 60-190 nmol in 10 ml culture volume), with smaller amounts of MHET and BHET detected throughout the observation period (Figure 3). The concentration of TPA steadily increased from day 3 to day 7, reaching a maximum of approximately 318±47.2 µM on day 7, indicating a slow but continuous degradation process over time. In contrast, strain UHH-Hm9b exhibited lower overall PET degradation activity. Nevertheless, UHPLC analysis confirmed its ability to release TPA from biofilms on PET foil at µM level. Interestingly, TPA concentration was highest on day 3 (41.6±13.2 µM in concentrated samples or 2.5±0.8 µM in original samples, corresponds to 25±8 nmol in 10 ml culture volume) but subsequently decreased to approximately 12±3.8 µM (7.2±2.2 nmol in 10 ml culture volume) on days 7 (Figure 3). Altogether these findings imply that UHH-5R5 and UHH-Hm9b form dense biofilms on non-treated PET foil and can release minute amounts of TPA under the tested conditions in the laboratory.

**Figure 3.**
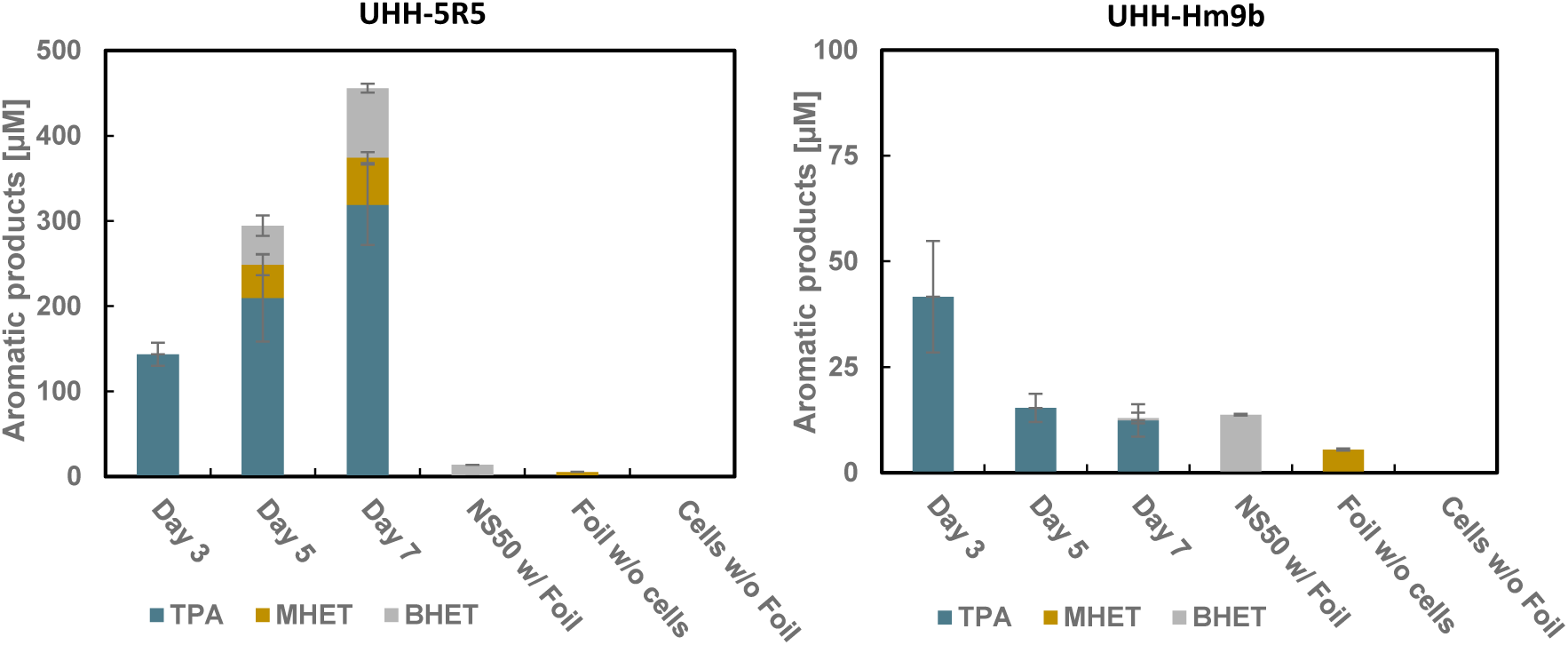
PET degradation products observed in supernatants from UHH-5R5 and UHH-Hm9b biofilms grown on PET film. Supernatants were collected after 3, 5, and 7 days and the degradation products were quantified using UHPLC as detailed in the Material and Methods section. Negative control samples were taken after 7 days of incubation. Data are normalized and corrected relative to *E. coli* DH5α growing on PET foil. PET degradation products in supernatants from UHH-5R5 and UHH-Hm9b biofilms grown on PET film. Values are mean of three biologically independent replicates, with error bars representing standard deviations.

### Genome sequencing identifies two novel dienelactone hydrolases involved in PET degradation

To identify the possible genes and enzymes linked to the observed hydrolytic activities, The genome of both isolates was sequenced. The draft genomes of both isolates were deposited at IMG under accession numbers 294449 and 294450. The genome assemblies implied a size of 10.2 Mbp for *Maribacter dokdonensis* UHH-5R5 and 6.1 Mbps for A*renibacter palladensis* UHH-Hm9b. A detailed analysis of both draft genomes and using the previously published HMM search motifs (21) for PETases revealed a single hit (e-value <1e-5) for possible PETases in each strain. The potential PET-acting esterase in Arenibacter corresponded to the predicted ORFs Ga0596863_004_127947_129248 and in Maribacter the ORF Ga0596861_0008_300124_301428 was identified as potential PETase (Figure 4A). The potential PET-esterase predicted in the two isolates were designated PET93 and PET94, respectively. Both hydrolase coding genes were not part of conserved regions on the bacterial chromosomes, and only a few other strains were observed with similar gene neighborhoods in the genomes available in IMG. None of the predicted genes were part of an operon, and the flanking genes appeared to be transcribed in the opposite direction. For both predicted genes putative promoter sequences were identified with a SD sequence 5-6 bp upstream of the translational start point.

**Figure 4.**
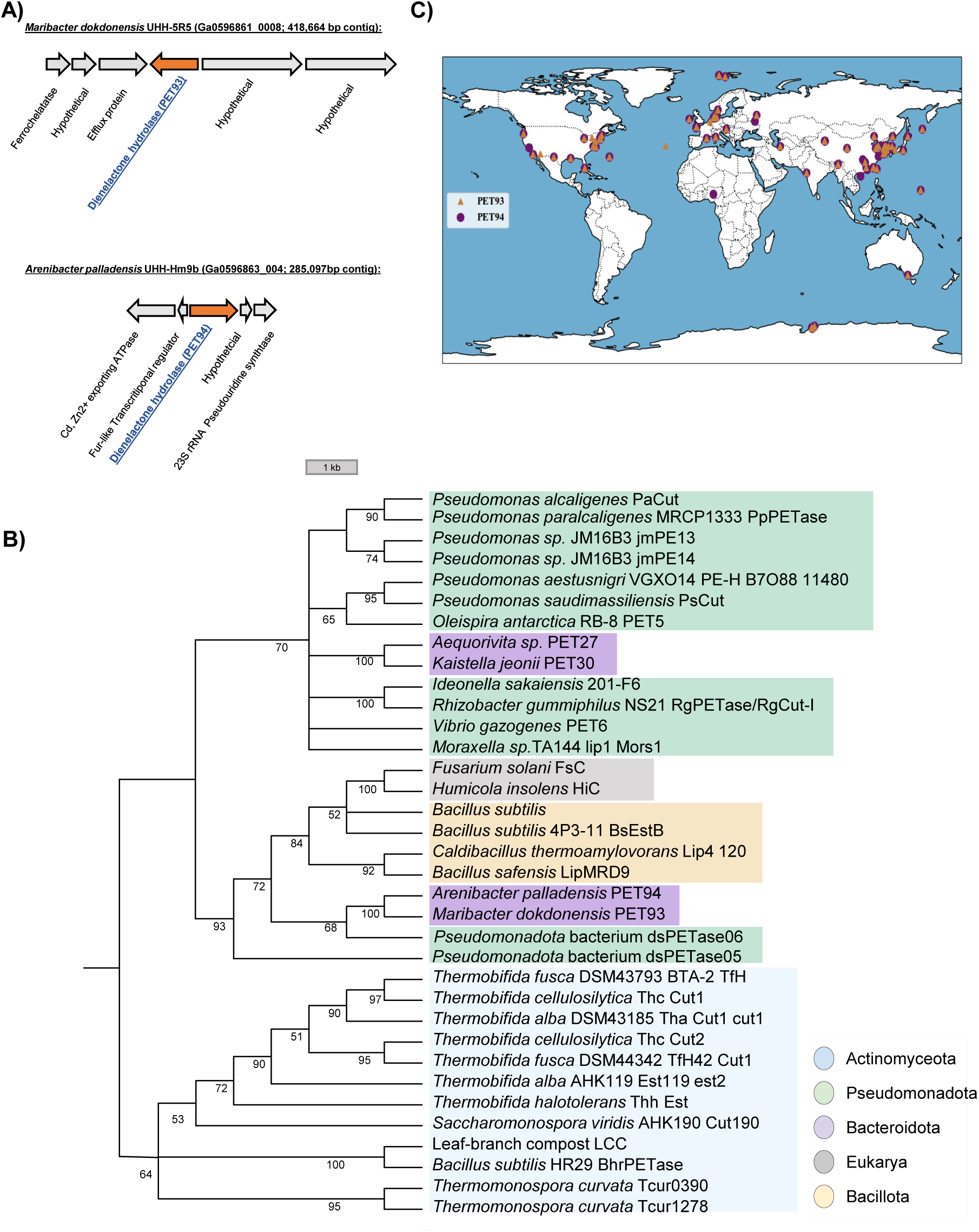
**A)** Genetic context of the dienelactone hydrolases PET93 and PET94 located in the Bacteroidota strains *Maribacter dokdonensis* UHH-5R5 and *Arenibacter palladensis* UHH-Hm9b, respectively; **B)** Neighbor-joining trees showing the phylogenetic relationship of PET93 and PET94 with other well-characterized enzymes. The tree was constructed using MEGAX and the number displayed at nodes are the bootstrap values based on 1,000 replications; **C)** Global distribution of PET93 and PET94 homologs. The corresponding contigs containing the two hydrolases are available for download from IMG using the indicated accession numbers. Enzyme entries were retrieved from the PAZy database.

The predicted esterase PET93 from *Maribacter dokdonensis* consisted of 434 amino acids with a possible secretion signal cleaving site identified between position 27 and 28 (VNA-QT) using the Sec/SPI secretion system. Similarly, the predicted protein PET94 derived from *Arenibacter palladensis* consisted of 433 amino acids and an N-terminal cleavage site between positions 25 and 26 (LNA-QT). Both amino acid sequences of the enzymes were highly similar (>99%) to precited DLHs in closely related species either affiliated with the genus Maribacter (PET93) or Arenibacter (PET94) in the NCBI database. However, their overall amino acid similarity to known PETases from the phylum of the Bacteroidota was relatively low with less than 50% identity (Table 2). Structural analysis also revealed a low similarity between the *Ideonella sakaiensis (Is)* PETase (*Is*PETase, PDB: 6EQE) structure and modeled structures of PET93 and PET94. Notably, the predicted active sites and binding sites of PET93 and PET94 were identical in their amino acid sequence to those of *Is*PETase and the leaf-branch compost cutinase (LCC, PDB: 4EBO) (Figure 7 & Table 2).

**Table 2:**
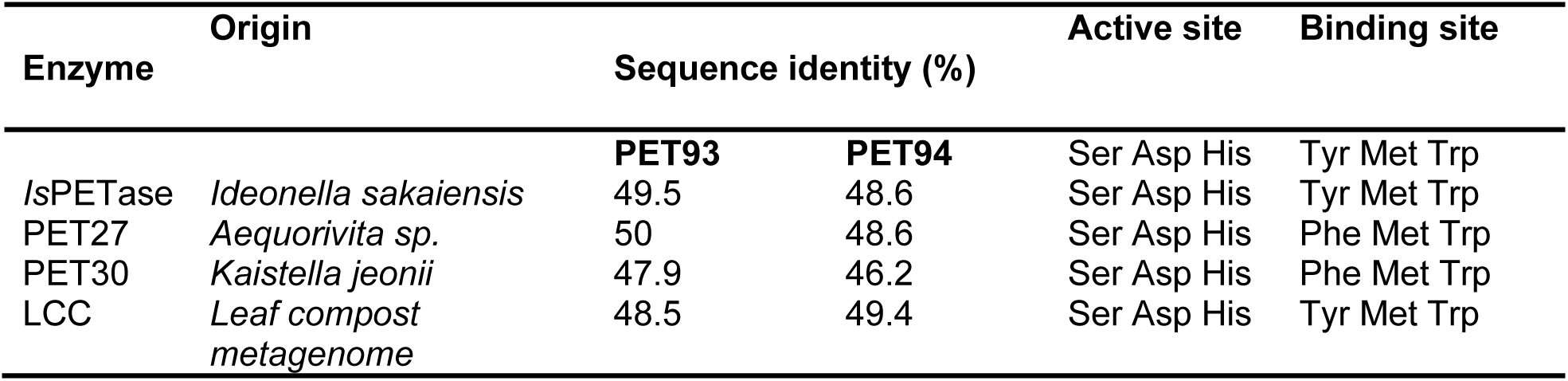
Sequence similarities generated for PET93 and PET94 against functionally verified PET hydrolases. Active site and PET binding motifs of PET93 and PET94 with selected and known PETases.

To evaluate the evolutionary relationships among bacteroidetal PET-hydrolyzing enzymes, including PET93, PET94 and other known PETases, a phylogenetic analysis was conducted using published and functionally verified PETases (Figure 4B). The resulting consensus tree revealed that putative Bacteroidota PET-degrading hydrolases formed a distinct subcluster. Interestingly, PET93 and PET94 were not grouped within the subcluster that included the previously known Bacteroidotal enzymes PET27 and PET30. They differed also with respect to the secretion signal. While PET93 and PET94 are secreted enzymes and code for a predicted N-terminally secretion signal, no PorC-like and C-terminal secretion signal was observed as it had been described for PET27 and PET30 previously.

### Partial biochemical characterization of potential PET93 and PET94

To further characterize both proteins and finally verify their PET-degrading activities, the gene sequences obtained by HMM were amplified from the corresponding organisms, cloned into *E. coli* DH5α *and* overexpressed in *E. coli* BL21(DE3). The transformed cells produced 48.1 and 45.6 kDa proteins when induced with 1mM IPTG. For both the C-terminal 6x histidine tagged proteins, we obtained recombinant proteins using Ni-NTA purification protocols with relatively high purity and activity on BHET plates (Figure 5A). We first characterized the activity of the two recombinant enzymes using *p*NP-assay. A substrate spectrum was recorded with *p*NP-esters which have acyl chain lengths of 4-18 carbon atoms. PET93 and PET94 revealed a narrow spectrum of substrates they hydrolyzed. The substrate specificity tests clearly showed that PET93 and PET94 both are capable of hydrolyzing ester bonds of a chain length of 6-10 carbon atoms but exhibit highest activity to hydrolyze a chain length of C8 *p*NP-octanoate (Figure 5B). The hydrolyzing activity of both enzymes was also accessed on the temperature range from 4 to 90°C. Both enzymes were shown to perform best at 40°C and interestingly they were able to act at 4°C (Figure 5C), implying a possible cold adaptation.

**Figure 5.**
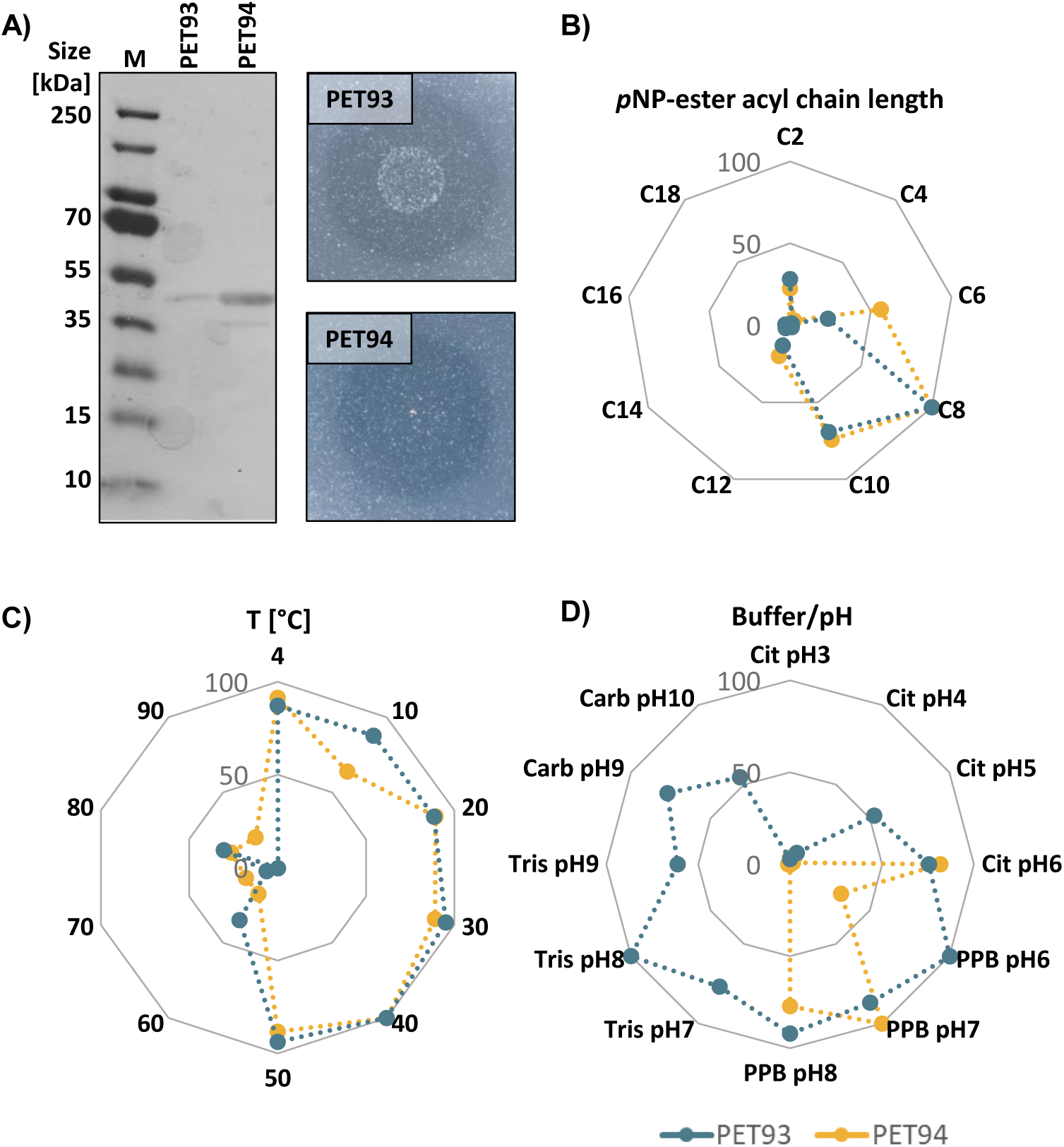
Purification and biochemical characterization of the dienelactone hydrolases PET93 and PET94. Purification of the His_6_-tagged dienelactone hydrolase PET93 (48.1 kDa) and PET94 (45.6 kDa) from *Maribacter dokdonensis* UHH-5R5 and *Arenibacter palladensis* UHH-Hm9b, respectively, and protein activity on BHET-containing plates **(A).** Substrate preference was tested with *p*NP-acetate (C2) to *p*NP-stearate (C18); **B).** Temperature **(B),** pH **(C)** and buffer optimum **(D)** were tested with *p*NP-octanoate C8. The assays were conducted using 0.1M buffers at pH 3-10 (pH 3-6, citrate buffer (Cit); pH 6-8, potassium phosphate buffer (PPB); pH 7-9, Tris buffer; pH 9-10, carbonate-bicarbonate buffer (Carb). All assays were carried out in triplicate at 37 °C for both enzymes. Data represent the mean values of three independent replicates, with standard deviation (SD) ≤ 10% for all measurements.

Buffers with pH between 3 and 10 were used to discover the optimal pH conditions for PET93 and PET94. With C8 *p*NP substrate, it was shown that PET93 was most active in 0.1M citrate buffer pH 6 and in 0.1M potassium phosphate buffer pH 7 while PET 94 was highly active in potassium phosphate buffer pH 6-8. PET93 lost its activity nearly completely in the citrate buffer at pH below 5, Tris buffer at pH values 7-9, and Carb buffer at pH values 9-10 (Figure 5D). Further it is shown that PET94 was almost inactivated under acidic conditions pH<4.

### PET93 and PET94 activity towards MHET, BHET and PET

Given that PET93 and PET94 are most active at 30-40°C in pH 7 potassium phosphate buffer, the recombinant enzymes were initially assayed with substrates MHET (1 mM) and BHET (5 mM). Therefore, UHPLC analysis was performed to identify the PET degradation product TPA (Table 3). Both enzymes can hydrolyze BHET and MHET to TPA and 0.2 mg mL^-1^ of PET93 released an average of 2642.8 ± 46.9 µM TPA from BHET and 776.3 ± 29.5 TPA µM TPA from MHET after 24 hours in 200 µl reaction volume. Under the same conditions, PET94 (0.2 mg mL^-1^) also released a similar amount of 2659.8 ± 99.6 and 684.5 ± 49.1 µM TPA when incubated with BHET and MHET, respectively.

**Table 3:** Recombinant and purified PET 93 and PET94 enzymatic hydrolysis of MHET (mono(2-hydroxyethyl) terephthalate), BHET (bis(2-hydroxyethyl) terephthalate) and PET (polyethylene terephthalate) with their final degradation product TPA (terephthalic acid). The concentration of the products was determined by UHPCL. The samples were incubated at 37^0^C for x hours with continuous shaking at 200 rpm and were analyzed in triplicates. BHET and MHET were added at 5 mM and 1 mM concentrations, respectively. *(*): PET substrates were pretreated with UV light for 1 week as indicated in the Material and Methods section*.

| Substrate added | Treatments | BHET detected (μM) | MHET detected (μM) | TPA detected (μM) |
| --- | --- | --- | --- | --- |
|  |  | Mean ± SD | Mean ± SD | Mean ± SD |
| BHET | Control | 2244 ± 38.4 | 286.6 ± 11.5 | n.d |
|  | PET93 | n.d | n.d | 2642.8 ± 46.9 |
|  | PET94 | n.d | n.d | 2659.8 ± 99.6 |
| MHET | Control | n.d | 736 ± 31.5 | 20.1 ± 0.73 |
|  | PET93 | n.d | n.d | 776.3 ± 29.5 |
|  | PET94 | n.d | n.d | 684.5 ± 49.1 |
| PET Foil* | PET93 | n.d | n.d | 29.8 ± 1.76 |
|  | PET94 | n.d | n.d | 37.3 ± 11.5 |
| PET Powder* | PET93 | n.d | n.d | 40.4 ± 18.67 |
|  | PET94 | n.d | n.d | 54.7 ± 21.9 |

n.d: not detected

Notably, when non-treated PET foil or powder was used as substrate for these two enzymes at pH 7 and 37°C, no TPA was detected by UHPLC after 5 days incubation. However, both enzymes were able to degrade UV-treated PET. After 5 days incubation with UV-treated PET powder, reasonable levels of PET degradation product were observed in a 200 μl reaction volume by UHPLC analyses, reaching 40.4 ± 18.67 µM and 54.7 ± 21.9 µM TPA released by PET 93 and PET94 (corresponds to 8.08 ± 3.73 nmol and 10.94 ± 4.38 nmol), respectively. However, only about 29.8 ± 1.76 and 37.3 ± 11.5 µM TPA by PET93 and PET94 (corresponds to 5.96 ± 0.35 nmol and 7.46 ± 2.30 nmol) was detected under the same conditions with UV-treated foil (Table 3). This suggests that UV probably helps promote the initial breakdown of PET.

Since we observed that the recombinant proteins had relatively low turnover rates in the µM range, we used the recently published *Comamonas thiooxidans (C. thiooxidans)* S23 reporter strains (Dierkes RF (2013) that is able to detect nM concentrations of TPA in further tests. Using this reporter strain (ReporTPA_UHH04, Supplementary Table S1) we were able to detect TPA release on PET foil incubated with recombinant PET93 and PET94 (Figure 6A). Controls incubated with BSA showed no fluorescence (Figure 6B). Altogether the data implied that the recombinant enzymes PET93 and PET94 are both active on PET albeit at relatively low levels.

**Figure 6.**
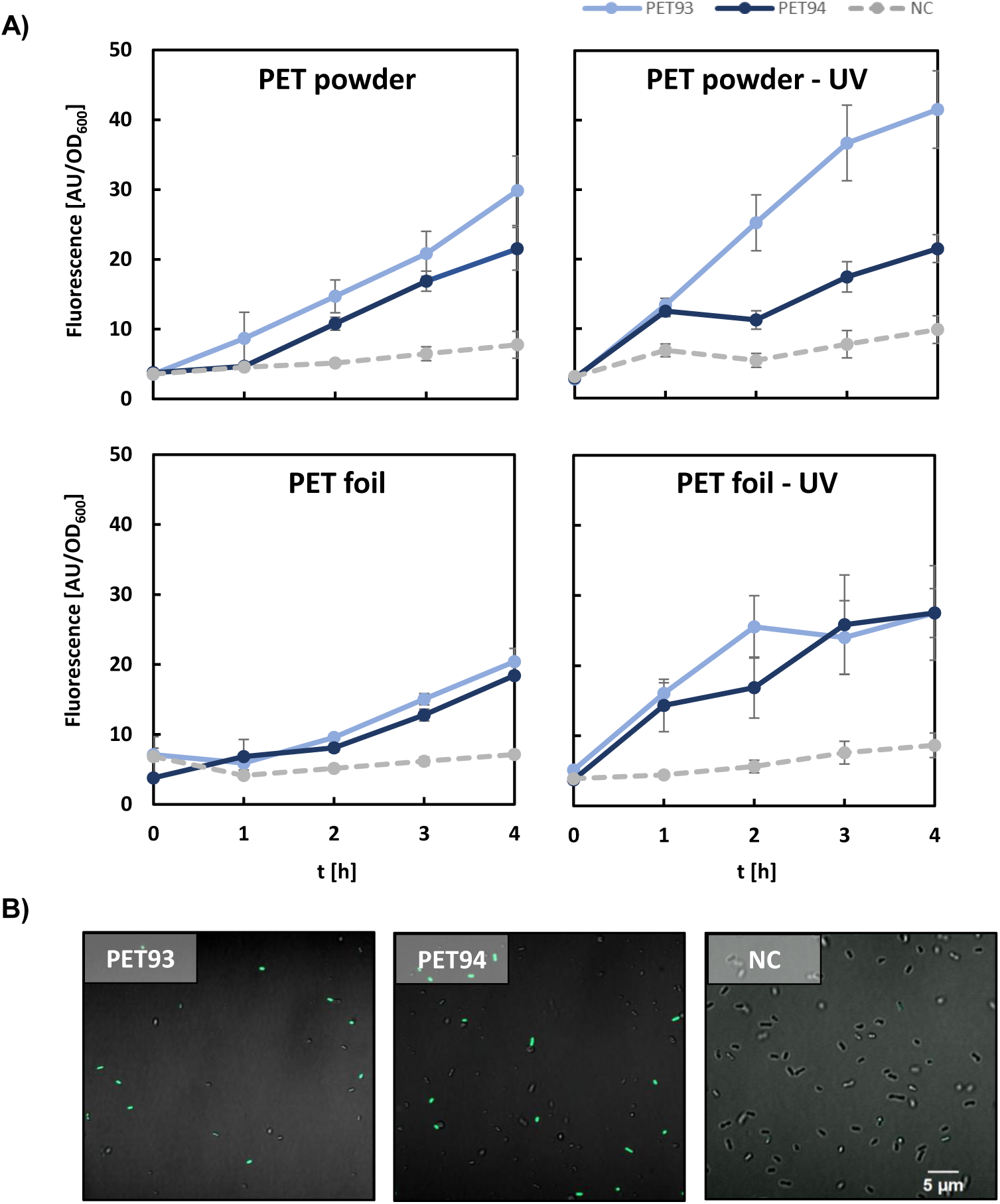
Detection of Enzymatic TPA Release from Untreated and UV-Treated PET Powders and Foils by *Comamonas thiooxidans* UHH04. Fluoresence response of *C. thiooxidans* UHH04 reporter cells (Dierkes, 2022) to supernatants from incubations of PET93 and PET94 with PET powder and foil, both untreated and UV-treated **(A).** Respectively 0.1 mg mL^-1^ of each enzyme and BSA as a negative control (NC) were incubacted with PET for 5 d in 0.1 M potassium phosphate buffer pH 7 before the addition of the UHH04 reporter to the supernatant. Fluorescence signals of sfGFP were normalized to the absorbance of the reporter cells at 600 nm. Data points represent mean values of 6 measurements, with standard deviation indicated by error bars. CLSM images of UHH04 reporter cells incubated with supernatants from the enzymatic reactions of PET93, PET94, and BSA (NC) on PET **(B).** The sfGFP fluorescence channel intensity was uniformly set for all images to allow for comparison.

**Figure 7.**
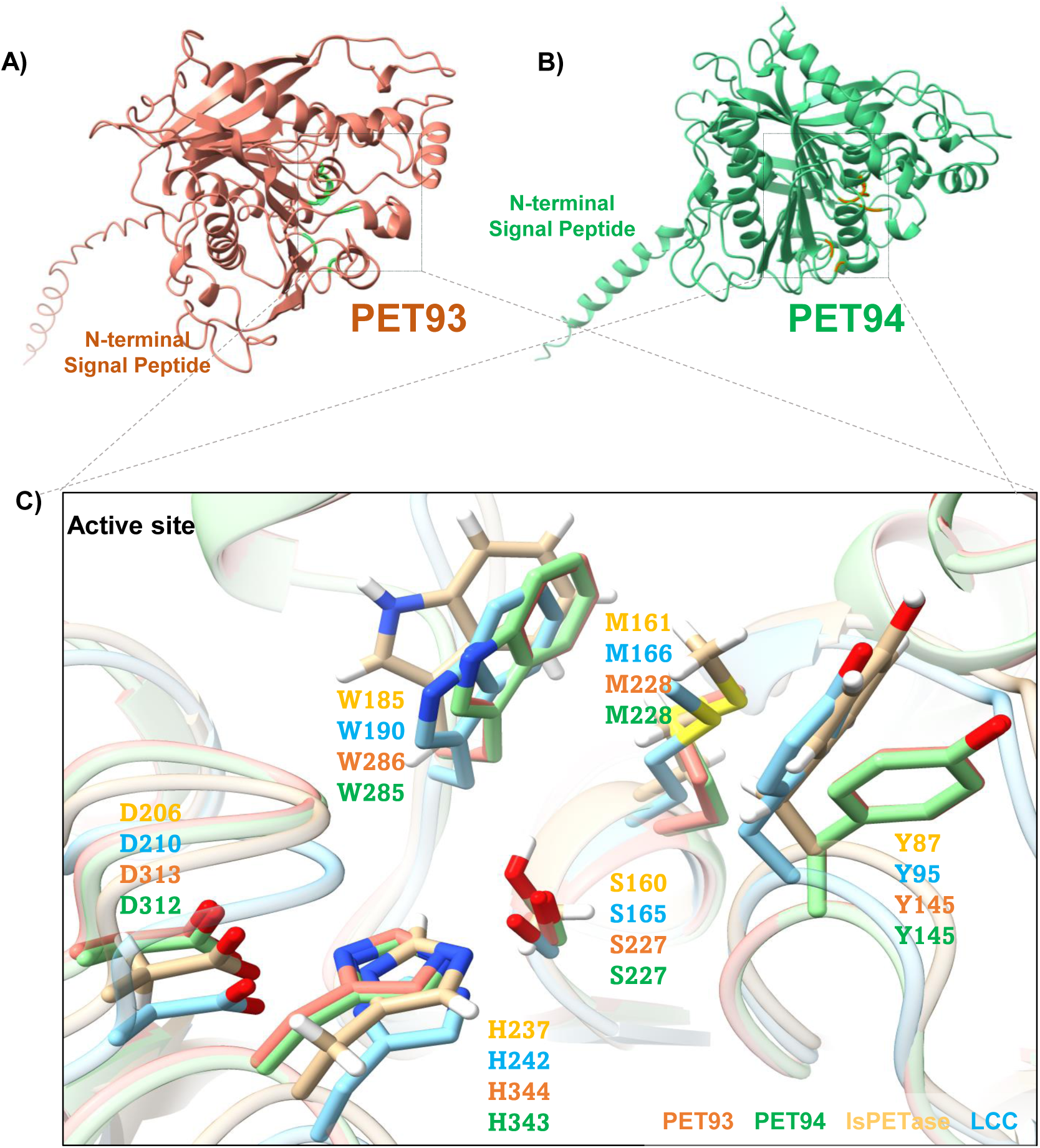
Structural model of PET-hydrolyzing enzymes from Bacteroidota. **A)** Overall structure of PET93 including active site and potential substrate binding site. **B)** Overall structure model of PET94 including active site and potential substrate binding site. **C)** Comparison of active site residues. All four enzymes PET93 (orange), PET94 (green), *Is*PETase (yellow, PDB 6EQE) and LCC (light blue, PDB 4EBO) have the typical residues of Ser-hydrolases at the catalytically active positions (Ser, His and Asp) and also have the same amino acids associated with PET-binding. The residues of *Is*PETase and LCC are indicated in light orange and light blue, respectively.

### Global distribution of PET93 and PET94

We further analyzed the global distribution of PET93 and PET94 and their homologs. It is notable that we were able to identify homologs in more than 250 currently available genome sequences of other closely related strains from publicly available databases IMG/MER (threshold of 50% sequence identity and over 80% coverage) (22, 23). Analysis of their occurrence and frequency of PET93 and PET94 raised the question of to what extent these enzymes could impact plastic degradation in the natural environment. For a focused view, we selected 99 homologs assigned to the Bacteroidota phylum in our global search to construct a global distribution map. Interestingly these homologs originated from a broad range of countries and regions (Supplementary Table S3), suggesting that the Bacteroideotal-derived enzymes PET93 and PET94 are widespread and may play a significant role in the global degradation of PET in nature. (Figure 4C).

## DISCUSSION

Recent studies showed that bacteria affiliated with the genera Maribacter and Arenibacter within the phylum of the Bacteroidota are globally distributed and play a significant role in the marine carbon cycle (24). They commonly inhabit marine sediments, coastal environments, and algae-associated habitats (25–30). Both genera demonstrate a broad and efficient capacity to degrade complex polysaccharides, playing a crucial role in organic matter cycling and microbial interactions in marine ecosystems (31–34). Notably, the occurrence of microbial species affiliated with both genera in the plastisphere has been described recently (35, 36). While it is well-known that bacteria in the plastisphere usually form biofilms on micro- or nanoparticles, it is commonly accepted that they mostly do not degrade the polymer (5, 37–39). However, it is assumed that they do primarily feed on the additives contained in the polymers and /or simply use the surface to attach and form biofilms (5, 40, 41).

While microorganisms in the phylum of the Bacteroidota are well known for their very diverse set of enzymes involved in polysaccharide hydrolysis, only a few species have been linked to plastic (i.e.PET) degradation to date (42). An initial HMM-based screen of metagenomic datasets has revealed that the Bacteroidota harbor considerable potential as yet underexplored sources of PET-degrading enzymes, particularly in the marine environment (21). The newly discovered enzymes PET93 and PET94 in this study are both promiscuous (E.C. 3.1.1.45). While the native substrate of both enzymes remains currently unknown, our data showed a significant but slow turnover of PET (Figure 3 and 6). Both enzymes differed largely in their structure from the previously published T9SS-dependent esterases involved in PET degradation. In fact, they represent novel scaffolds and imply that the phylum of the Bacteroidota harbors a diverse set of promiscuous enzymes to degrade these man-made polymers.

Notably, homologs of PET93 and PET94 were found globally in genome data sets covering a wide range of climate zones (Figure 4C). It is likely due to the ability of Bacteroidota to decompose a wide range of biopolymers, including cellulose, algal polysaccharide (e.g. laminarin, alginate, xylan) and other complex carbohydrates (43–45). Similar to other Bacteroidotal-enzymes PET27 and PET30 (42), our enzymes demonstrated PET-hydrolyzing activity at temperature as low as 4°C, suggesting a potential role in slow, long-term degradation of PET microparticles in cold environments. While these data do not prove that PET93 and 94 are active in nature, it is likely that they are secreted and will be involved in enzymatic PET degradation. Recently we showed that in *Vibrio gazogenes* a PETase (PET6) was expressed constitutively at a low but significant level under various growth conditions (46). Assuming that in Maribacter and Arenibacter similar regulatory pathways may exist, it is likely that PET93 and 94 are also expressed at low levels under biofilm and planktonic growth conditions.

Although the catalytic efficiency of PET93 and PET94 towards PET do not meet the level of industrial applications, their ecological significance should not be underestimated as they are derived from globally occurring microorganisms. Their global presence highlights that these microbes may play an important but not yet fully understood role in natural bioremediation of marine plastic pollution. Thus, future work will have to explore the role of the secreted DLHs PET93 and PET94 in their native environment.

## MATERIALS AND METHODS

### Enrichment, isolation and identification of Bacteroidota strains

Microbial communities from marine and aquatic environments were enriched in 100ml R2A or BMB media with 1g of PET powder. Enrichment cultures were then incubated at 22°C and 28°C under continuous shaking conditions. After enrichment, 100µl of the cultures were spread on the R2A/BMB agar plate. The 16S rDNA gene of the colonies was amplified using standard 16S primers. The amplified fragments were Sanger sequenced afterwards at Eurofins (Elsberg, Germany). The colonies were then screened for their hydrolysis activity on tributyrin (TBT), polycaprolactone (PCL) and bis-(2-hydroxyethyl) terephthalate (BHET). The strains with hydrolytic activities observed towards TBT, PCL or BHET were later selected for whole-genome sequencing. Genomic DNA (gDNA) were extracted with the NucleoSpin® Microbial DNA Kit from MN (Düren, Germany) from 5 mL cultures and then sequenced using IIIumina NextSeq 500 sequencing method at Eurofins (Germany).

Bacterial strains and plasmids used in the study are listed in Table S1. *Escherichia coli* was grown in LB medium (1% tryptone/peptone, 0.5% yeast extract, 1% NaCl) supplemented with appropriate antibiotics at 250 rpm in flasks under aerobic conditions at 37°C for 20 hours.

### Data availability and bioinformatic analysis

The genome of isolates UHH-5R5 and UHH-Hm9b that originated from marine aquaculture were submitted to IMG.gov under the submission IDs 294449 and 294450, respectively. The sequence reads were assembled using SPAdes (v.3.15.0) (47) and initial annotated with Prokka (v 1.14.6) (48). To identify putative PET esterases within the *Bacteroidetes* genome datasets, a profile Hidden Markov Model (HMM) was constructed based on the known, functionally tested enzymes (49). The HMM analysis identified PET93 and PET94 as homologues of known PET degrading enzymes, which were subsequently investigated further.

Nucleotide and amino acid sequences of putative PET esterases were obtained from genomic data of isolates UHH-5R5 and UHH-Hm9b from IMG. Sequence data were processed and analyzed using Snapgene (GSL Biotech LLC, San Diego, CA, USA). Conserved domains in the protein sequences were identified using CD-search (50). A phylogenetic tree was constructed with MEGA-X, employing maximum bootstrap of 1000 for enhanced accuracy (51). Structural information was retrieved from the RCSB-PDB database (52), and protein structures were predicted using AlphaFold2 with default parameters (53). The 3D protein models were visualized in UCSF Chimera, further the structural alignment with homologous protein were generated with Chimera MatchMaker tool (54). SignalP 5.0 server was used to predict the native signal peptide sequences (55).

### Biofilms on PET surface: growth and degradation product analysis

Precultures of UHH-5R5 and UHH-Hm9b were inoculated in BMB media at 28°C with continuous shaking at 130 rpm. The cultures were then diluted to an OD_600_ of 0.05 and then 5 ml of each diluted cultures were transferred to 6-well plate (Nunc cell culture plate, catalog no. 130184; Thermo Fisher Scientific, Waltham, MA). PET foil platelets (Ø 35 mm, amorphous PET film, Goodfellow GmbH, Bad Nauheim, Germany) were sterilized in Ethanol 70% for 10 mins and then added to each well, which were subsequently incubated at 28°C under 80 rpm shaking condition to facilitate biofilm formation. Supernatant was collected after 3, 5 and 7 days of incubation. Each 10 ml of supernatant in each sample was vacuum dried to a final volume of 600 µl.

The concentrated supernatants were further analyzed using UltiMate™ 3000 UHPLC system (Thermo Fisher Scientific, Waltham, MA, USA). The Triart C18 column (YMC Europe GmbH, Dinslaken, Germany), 100 × 2.0 mm with 1.9 µm diameter was employed for separation. Isocratic elusion was performed with a mobile phase consisting of 20:80 (v/v) acetonitrile and water (acidified with 0.1% v/v trifluoroacetic acid) at a flowrate of 0.4 mL min^-1^. 50 µl of concentrated supernatant was mixed with 200 µl of acetonitrile (acidified with 1% vol trifluoroacetic acid), followed by centrifugation at 10,000 × g for 3 min. A 200 µl aliquot of the mixture was then diluted with 600 µl water. Each 15 µl of sample was then injected for each measurement. As negative controls, PET foils are incubated in media without bacterial cells and only medium was incubated under the same conditions as *E. coli* Dh5α. These samples were taken at the same time point. All experiments were performed with 3 biological replicates.

### Imaging analysis of biofilms on PET foil platelets

For observation of biofilm formation of Bacteroidetes isolates UHH-5R5 and UHH-Hm9b on plastic surface, the cells were grown in BMB medium at 28°C with 130 rpm shaking until reaching an optical cell density OD_600_ of 1. These starter cultures were then diluted to the OD_600_ of 0.05 in fresh BMB medium. The cultures were incubated with PET foils at 28°C with 80 rpm shaking. After incubation, foil was washed three times with 1x PBS buffer and placed to μ-Slide 8-well plates (ibiTreat. catalog no. 80826, ibidi USA, Inc., Fitchburg, Wisconsin). Cells were stained using 100 µL of the LIVE/DEAD® BacLight™ Bacterial Viability Kit (Thermo Scientific). Cells were then analyzed using the Axio Observer Z1/7, LSM 800 confocal microscope equipped with an objective C-Apochromat 63x/1.2 W Korr UV VisIR (Carl Zeiss Microscopy GmbH, Jena, Germany), utilizing the SYTO-9 channel (emission wavelength: 528/20 nm) and the PI channel (emission wavelength: 645/20 nm). For the analysis of the CLSM images the ZEN software was used (version 2.3. Carl Zeiss Microscopy GmbH, Jena, Germany). For each sample at least three different positions were observed and one representative CLSM image was chosen.

### Heterologous expression of recombinant putative enzymes

PET93 and PET94 were amplified from genomic DNA of UHH-5R5 and UHH-Hm9b and cloned into pET21a (+) vector. The constructs were sequenced at Microsynth Seqlab GmbH (Goettingen, Germany) and compared to the original sequence to check for the correctness. Sequences coding mature PET94 (sequence devoid of the signal peptide) and PET93 protein were heterologous expressed in *E. coli* BL21 (DE3) using *β*-D-1-thiogalactopyranoside (IPTG) induction. Cultures were incubated aerobically in LB medium with ampicillin 100 μg/ml at 37°C. When OD_600_ reached 0.7-0.8, the expressions were induced with IPTG 1mM, followed by incubation at 22°C and 17°C for 20 hours. Cells were harvested and lysed with pressure using a French press. Afterwards the proteins with C-terminal 6x histidine tag were purified via Nickel-ion affinity chromatography using Ni-NTA agarose (Qiagen, Hilden, Germany) and analyzed by SDS-PAGE. The elution buffer was exchanged against 0.1 mM potassium phosphate buffer pH 7.0 in a 30 kDa Amicon Tube (GE Health Care, Solingen, Germany).

### Plate-based activity assay and partial biochemical characterization of PET93 and PET94

For activity tests, purified recombinant proteins were utilized. Agar plates were prepared containing 10mM bis-(2-hydroxyethyl) terephthalate (BHET) and 500 mg L^-1^ polycaprolactone (PCL). 10 µl of eluate from protein purification were spotted onto the plates to observe the halo formation.

For the *p*NP-assay, unless otherwise specified, 0.1–1 µg of the enzyme was added to a substrate solution containing 190µl of 0.1 M potassium phosphate (pH 7-8) and 10 µl of 0.1mM pNP-substrates dissolved in isopropanol. The reaction was terminated after 10 minutes by adding 200mM of Na₂CO₃. The samples were then centrifuged at 4°C and 13,000 rpm for 3 minutes. Various *p*NP esters substrates with chain lengths of C4, C6, C8, C10, C12, C14, C16 and C18 were tested. Enzyme activity was indicated by the color change from colorless to yellow, with the absorbance measured at 405nm in a plate reader (Biotek, Winooski, USA). All measurements were performed in triplicate. The optimal temperature for enzyme activity was evaluated between a range of 10 to 90°C. Additionally, the effect of pH on enzyme activity was accessed using citrate phosphate buffer (pH 3.0, 4.0, and 5.0), potassium phosphate buffer (pH 6.0, 7.0, and 8.0) and carbonate bicarbonate buffer (pH 9.2 and 10.2) with pNP-C8 as the substrate.

### UHPLC-based activity assay for PET, MHET and BHET degradation

Purified proteins at the concentration of 0.1mg mL^-1^ were incubated with MHET (1 mM), BHET (10 mM), PET powder (10 g mL^-1^), PET foil/powder and UV-treated PET foil/powder in 200 µl of potassium phosphate buffer (pH 7-8). After 24 hours of incubation at 37°C with MHET/BHET and 120 h with PET, the supernatant was filtered through 0.22 µm filter paper. The release of TPA was then analyzed using UHPLC and TPA reporter strain *C. thiooxidans* UHH04 (56). BSA incubated with PET substrates under the same conditions served as a negative control.

### *C. thiooxidans* S23 reporter strain preparation for TPA Assays

The protocol for this experiment was adapted from Dierkes’s original method (56). The *C. thiooxidans* S23 biosensor strains ReporTPA_UHH04 were incubated overnight at 130 rpm in 50ml LB medium containing 25 µg/mL chloramphenicol and additionally supplemented with 10 mM gluconate in an Erlenmeyer flask. Before performing the TPA assays, the OD600 of the cultures was measured, and an appropriate volume was centrifuged at 4,500 rcf and 4°C for 5 minutes. The resulting pellet was resuspended in 50 ml Wx medium containing 25 µg/mL chloramphenicol to achieve a final OD_600_ of 0.6. The resuspended cultures were incubated at 37°C and 130 rpm for 30 minutes before being added to the samples.

For standard assay, 100 µL of the sample was added to each well of a black-walled 96-well microtiter plate (ThermoFisher, Waltham, MA, USA) designed for fluorescence-based assays. An additional 100 µL of reporter cells UHH04, prepared as described above, was added to each sample well. The plate was incubated at 28°C on a Vibration Shaker 3023 (Gesellschaft für Labortechnik mbH, Burgwedel, Germany) at 150 rpm. Fluorescence and OD600 measurements were taken at intervals of 0.5 to 2 hours using a Synergy HT plate reader with Gen5 software (BioTek, Winooski, VT, USA).

### Fluorescence microscopy of *C. thiooxidans* S23 reporter strain

Microscopic imaging of reporter cells was performed with a confocal laser scanning microscope Axio Observer.Z1/7 LSM 800 (Carl Zeiss Microscopy GmbH, Jena, Germany) using Plan-Apochromat 100×/1.40 Oil DIC M27 objective. Image analysis and processing was carried out using ZEN software (Version 2.3, Carl Zeiss Microscopy GmbH).

### Global distribution of PET93 and PET94 homologs

The IMG/M scans for PET93 and PET94 homologs were completed on April 20, 2025. When available, Geo locations were used as provided on IMG. In case the data was missing, we attempted to retrieve Geo coordinates using details about isolation source/location/city/country on IMG database. The map illustrates both the frequency and geographical distribution of the homologs of dienelactone hydrolases in the strains UHH-5R5 and UHH-Hm9b was created using the Cartopy Python package (version 0.24.0) that is freely available on https://scitools.org.uk/cartopy. A similarity threshold of 50% was applied in homology searches. Only bacterial hits classified within the Bacteroidota phylum were included in the final dataset.

## ACKNOWLEDGEMENTS

This work was supported by a grant from German Federal Ministry of Education and Research - BMBF (Funding reference number 031B0846G).

## SUPPLEMENTARY TABLE LEGENDS

**Table S1:** Bacterial strains and plasmids used in this study.

**Table S2:** Quantitative analysis of biofilm morphology formed by the Bacteroidota isolates UHH-5R5 and UHH-Hm9b on PET foil over a 7-day incubation period using BiofilmQ. Representative data of *n = 3* independent biofilms for each treatment are presented.

**Table S3:** Homologs of PET93 and PET94 hydrolases from Bacteroidota isolates UHH-5R5 and UHH-Hm9b. Data were retrieved from the publicly available IMG/MER database, applying thresholds of ≥50% sequence identity and ≥80% sequence coverage.

## Notes

The authors declare no conflict of interest.

